# Does Infection Progression of *Mycobacterium abscessus* Depend on Sex, Age, and Mouse strain?

**DOI:** 10.64898/2026.03.25.714167

**Authors:** Maya Rima, Aurélie Chauffour, Régis Tournebize, Corentin Poignon, Thibaud Coradin, Alexandra Aubry, Nicolas Veziris

## Abstract

The lack of a reliable chronic murine model limits drugs evaluation against *Mycobacterium abscessus*. Models show discrepancies, especially regarding host factors (mouse strain, sex and age). Using beads-model, we compared BALB/cJRJ and C57BL/6NCrl across sexes and ages. BALB/cJRJ showed more sustained infection and lower variability, with no significant sex- or age-related differences. Considering these results and the higher prevalence of NTM pulmonary infections in female patients, 5-6 weeks-old female BALB/cJRJ are appropriate for *M. abscessus* beads-model.

---

*Mycobacterium abscessus* (*M. abscessus*), described as “clinical and antibiotic nightmare”, is a rapidly growing nontuberculous mycobacterium (NTM) responsible for opportunistic pulmonary infections, particularly in individuals with underlying lung diseases. Its intrinsic and acquired resistance to multiple antibiotic classes renders treatment highly challenging, highlighting the urgent need for novel anti*-M. abscessus* drugs (1). However, such research is limited by the lack of a reliable and reproductible murine model of a chronic *M. abscessus* infection suitable for preclinical testing (2). To date, none of the infection models described in the literature fully satisfies the criteria recommended for a robust model of chronic infection : use of an affordable immunocompetent mice, use of the reference *M. abscessus* strain ATCC 1997, lung bacterial burden decrease <0.5 log_10_ CFU over 28 days, bacterial loads >5 log_10_ CFU at treatment start and end, and lung pathology demonstrating cellular granulomas with intracellular and extracellular bacteria (2).

One approach that has shown promising results in the literature is the encapsulation of *M. abscesssus* in agar beads to reduce bacterial clearance from the lungs, which represents a major obstacle in the development of a chronic infection model. However, substantial heterogeneity exists among the reported beads-based models, particularly regarding the sex and age of the animals used, which may contribute to the variability observed in the infection progression (3-5).

In the first study investigating this strategy for *M. abscessus*, a persistent infection lasting up to 90 days in 8-10 weeks-old male C57BL/6N mice has been reported (3). The use of males was supported by findings from Yamamto *et al*., who showed that males are more susceptible than females to developing severe pulmonary mycobacterial infections (6). However, a subsequent study demonstrated a comparable persistent infection in 6-8 weeks-old female C57BL/6JN mice, indicating that *M. abscessus* infection is not preferentially established in male mice (4). In contrast, and further complicating the interpretation, Malcolm *et al*., reported rapid bacterial clearance in female C57BL/6J mice, with no detectable load in the lungs of 3 out of 5 mice at 50 days post-infection (5).

Given these discrepancies among published results, there is a clear need for a systematic comparative evaluation. The objective of this study was to assess comparatively the impact of 1) mouse strain, 2) sex, and 3) age on host susceptibility to *M. abscessus* infection.

Animal experiment was performed according to ethical guidelines and approved by the ethical committee (APAFIS #31933-2021040214468842 v11). Briefly, BALB/cJRJ (Janvier breeding center, Le Genest-Saint-Isle France) and C57BL/6NCrl (Charles River, Saint Germain Nuelles, France) mice were intratracheally inoculated with 50 µL of *M. abscessus* (ATCC 19977, rough variant) encapsulated in alginate beads (5.04 log_10_CFU/mouse) following anesthesia by intraperitoneal injection of ketamine (100 mg/kg) and xylazine (10 mg/kg) (S1, Table S2). Both females and males of different ages (5-6 weeks-old and 8-10 weeks-old) were included in this study to enable comparison with the literature. Bacterial load in the lungs of mice was quantified after 1-, 7-, 14-, and 30-days post-infection (dpi) after grinding the tissue in 2 mL of sterile water (Versol, Dutscher) using a gentleMACS™ dissociator. Bacteria were then plated on 7H11 agar medium (212203, BD) supplemented with 0.4% charcoal (C9157, Sigma-Aldrich) and antibiotics (Mycobacteria Selectatab, MS24, MastGroup) and incubated for 5 days at 30°C.

Results showed that, in BALB/cJRJ mice, the bacterial load followed a similar pattern in females and males across the tested age groups (Figure 1a). Bacterial burden remained relatively stable up to 14 dpi in all groups, except in 8-10 weeks-old females, which exhibited ∼ 1 log_10_CFU/lungs reduction at 14 dpi compared with 1 dpi (Table 1). By 30 dpi, bacterial load significantly decreased in all groups regardless of the sex of the mice, although a trend toward slower clearance was observed in males (2 to 2.2 log_10_CFU/lungs reduction between 1 and 30 dpi compared with 2.7 to 2.8 log_10_CFU/lungs reduction in females).

**Table 1.** *M. abscessus* bacterial load recorded in the lungs of BALB/cJRJ mice.

| BALB/cJrJ |  |  |  |  |  |
| --- | --- | --- | --- | --- | --- |
| Sex | Age (weeks) | Bacterial load in lungs (log<sub>10</sub> CFU/lungs ± SD) |  |  |  |
|  |  | 1 dpi | 7 dpi | 14 dpi | 30 dpi |
| Female | 5-6 | 5.62 ± 0.26 | 6.35 ± 0.35 | 5.02 ± 0.52 | 2.92 ± 0.33 |
|  | 8-10 | 5.68 ± 0.41 | 5.91 ± 0.27 | 4.50 ± 0.26 | 2.86 ± 0.57 |
| Male | 5-6 | 5.62 ± 0.35 | 6.74 ± 0.21 | 5.31 ± 0.37 | 3.46 ± 0.20 |
|  | 8-10 | 5.69 ± 0.46 | 6.06 ± 0.26 | 5.43 ± 0.33 | 3.67 ± 0.44 |

**Figure 1.**
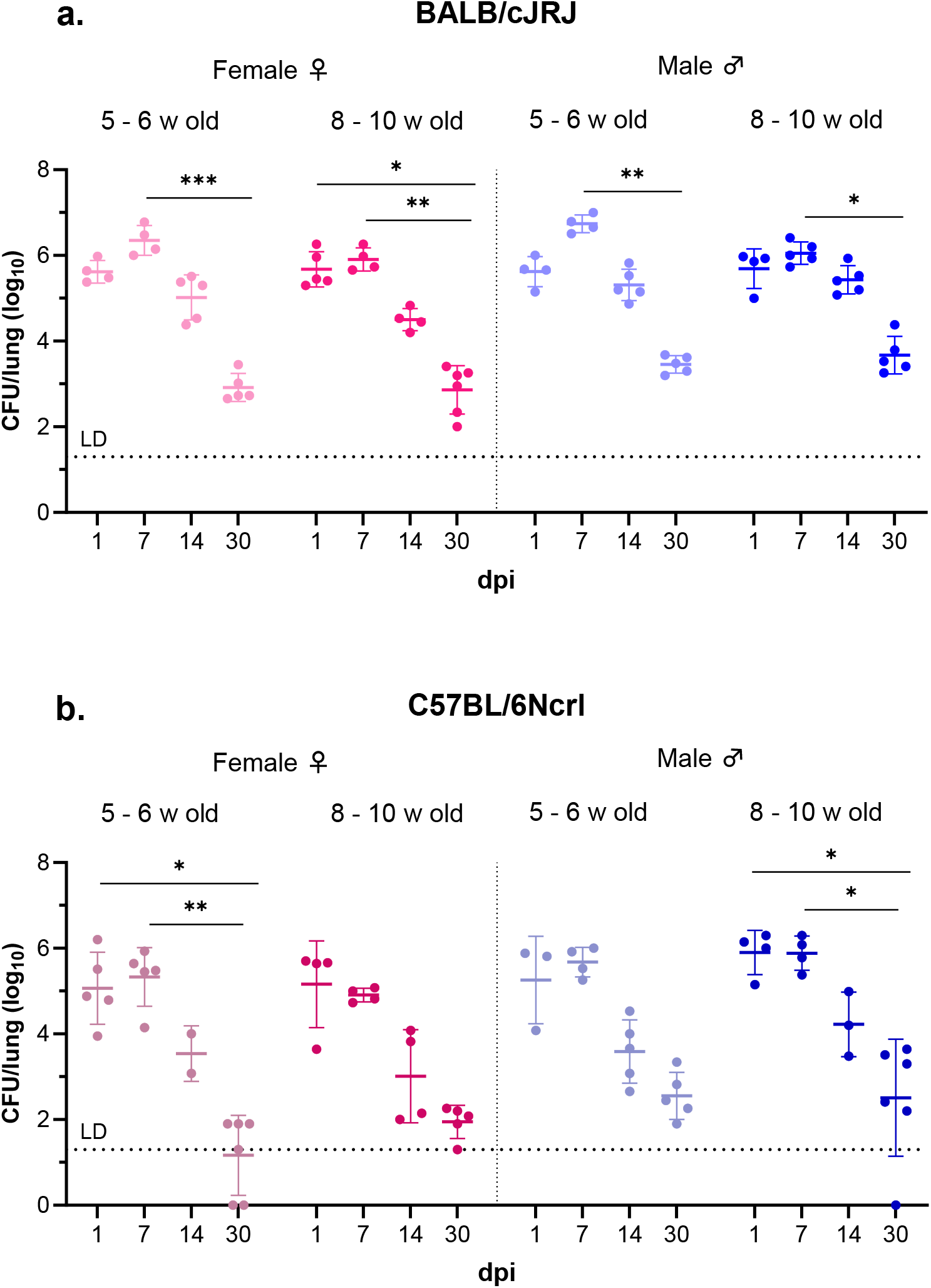
*M. abscessus* burden in the lungs of BALB/cJRJ (a) and C57BL/6Ncrl (b) mice of different sexes and ages, intratracheally infected with bacteria encapsulated in alginate beads (5.04 log_10_ CFU/mouse), determined at 1, 7, 14, and 30 dpi (LD: limit of detection). Statistically significant differences were determined by one-way ANOVA with Kruskal-Wallis test for multiple comparisons across all time points within the same sex and age group (*, *p-* value < 0.05; **, *p-*value < 0.01; ***, *p-*value < 0.001).

Regarding C57BL/6NCrl, bacterial clearance was faster than in BALB/cJRJ, with all groups showing a reduction of more than 1.5 log_10_CFU/lungs by 14 dpi (Figure 1b, Table 2). Compared to 1 dpi, by 30 dpi bacterial clearance was between 2.7 and 3.9 log_10_CFU/lungs in C57BL/6NCrl mice, whereas it ranged from 2 to 2.8 log_10_CFU/lungs in BALB/cJRJ mice. Surprisingly, CFU counts exhibited notably higher variability within groups compared with BALB/cJRJ mice, even though both mouse strains were inoculated on the same day and with the same inoculum. This variability may explain the differing findings observed across beads-based published models using this mouse strain (4, 5). As in BALB/cJRJ mice, we did not observe any significant age- or sex-related differences in C57BL/6NCrl mice.

**Table 2.** *M. abscessus* bacterial load recorded in the lungs of C57BL/6NCrl mice.

| C57BL/6NCrI |  |  |  |  |  |
| --- | --- | --- | --- | --- | --- |
| Sex | Age (weeks) | Bacterial load in lungs (log<sub>10</sub> CFU/lungs ± SD) |  |  |  |
|  |  | 1 dpi | 7 dpi | 14 dpi | 30 dpi |
| Female | 5-6 | 5.07 ± 0.84 | 5.33 ± 0.69 | 3.54 ± 0.65 | 1.17 ± 0.93 |
|  | 8-10 | 5.16 ± 1.01 | 4.91 ± 0.16 | 3.01 ± 1.09 | 1.95 ± 0.39 |
| Male | 5-6 | 5.26 ± 1.02 | 5.68 ± 0.35 | 3.59 ± 0.74 | 2.55 ± 0.55 |
|  | 8-10 | 5.90 ± 0.52 | 5.88 ± 0.4 | 4.22 ± 0.75 | 2.51 ± 1.36 |

Overall, in our beads model, BALB/c mice were more susceptible to *M. abscessus* than C57BL/6. This differential susceptibility of these mouse strains to mycobacterial infections remains controversial. Indeed, some studies report that BALB/c mice are more susceptible than C57BL/6 to mycobacterial infections (6-8). This difference is often attributed to immunological factors: BALB/c preferentially generate a Th2-type immune response, whereas C57BL/6 primarily develop a Th1-type immune response, which is known to be a protective response against intracellular pathogens, including *M. abscessus* (9, 10). Nevertheless, some studies did not observe differences between these two mouse strains in *M. abscessus* infection, highlighting the complexity of host-pathogen interactions in mycobacterial disease (11, 12).

Regarding the impact of the sex, we did not observe any significant differences between males and females, regardless the mouse strain. The influence of the sex on mycobacterial infections remains controversial in the literature. While some studies report no sex-related differences, other describe sex-dependent variations in infection outcomes (13-15). Indeed, an early study demonstrated that males BALB/c develop more severe *M. intracellulare* infection compared to females, which is associated with reduced phagocytic activity of peritoneal macrophages in males (16). The same group subsequently reported similar findings in distinct mouse strains infected with *M. marinum*, with the most pronounced sex-related differences observed in BALB/c compared with C57BL/6. They suggested that testosterone may contribute to the increased susceptibility observed in males (6).

Altogether, as the decrease in CFU was not significantly lower in males than in females, we recommend the use of female mice to optimize a *M. abscessus* chronic infection model for two main reasons: 1) NTM pulmonary infections in humans predominately affect female patients (17), and 2) female mice are less aggressive and therefore easier to house and handle, which is a key consideration when conducting experiments involving large numbers of mice.

In conclusion, in our beads model, 5-6 weeks old BALB/c mice exhibit a more consistent and prolonged *M. abscessus* infection than the other mouse groups evaluated, with females recommended given the absence of significant sex-related differences. Moreover, their low intra-group variability makes them suitable for evaluating the efficacy of novel drugs.

## Acknowledgments

This project has received funding from the Innovative Medicines Initiative 2 Joint Undertaking (JU) under grant agreement No [853932]. The JU receives support from the European Union’s Horizon 2020 research and innovation programme and EFPIA.

## Supplementary materials

**S1: Preparation of *M. abscessus* suspension encapsulated in alginate beads**

A fresh *M. abscessus* liquid culture was grown in 7H9 medium (217310, BD) supplemented with 10% OADC (211886, BD), and 0.05% of Tween 80 (2002-C, Euromedex) for 4-5 days (30°C, 80 rpm). The encapsulation of bacteria in alginate beads was performed using a Nisco encapsulation system (VARJ30, Nisco Engineering AG, Zurish, Switzerland). The prepared bacterial suspension (OD_600nm_ = 0.6-0.7) was mixed with a 1% alginate solution (Alginic acid sodium salt, 71238, Sigma-Aldrich) prepared in 0.9% NaCl (S/3160/60, Fisher) at a 1:2 bacteria-to-alginate volume ratio. The mixture was then injected into the Nisco system. Bacterial encapsulation was carried out under a pressure of 40 mbar and a flow rate of 0.2 mL/min. Beads carrying bacteria were recovered in a slightly agitated gelling bath containing 0.1 M CaCl_2_ prepared in 0.1 M Tris/HCl solution (T1503, Sigma-Aldrich) and supplemented with 0.05% of Tween 20 (P1379, Sigma-Aldrich) to prevent beads aggregation. After one hour of stabilization in the gelling bath, the beads were washed twice by centrifugation (1500 rpm, 7 min) using a washing buffer containing 0.1 M CaCl_2_ in 0.9% NaCl. Since beads size is crucial and small beads can be easily cleared by mice, the prepared encapsulated bacterial suspension, diluted in 20 mL of the washing buffer, was filtered using a cell strainer (ClearLine Cell strainers 100 µm, 141380C, Dutscher) in order to discard beads smaller than 100 µm. Beads with a size of 100 µm or larger were collected by rinsing the inverted cell strainer with the washing buffer. Encapsulated bacteria were enumerated on 7H11 agar medium (5 days, 30°C).

**Table S2.**
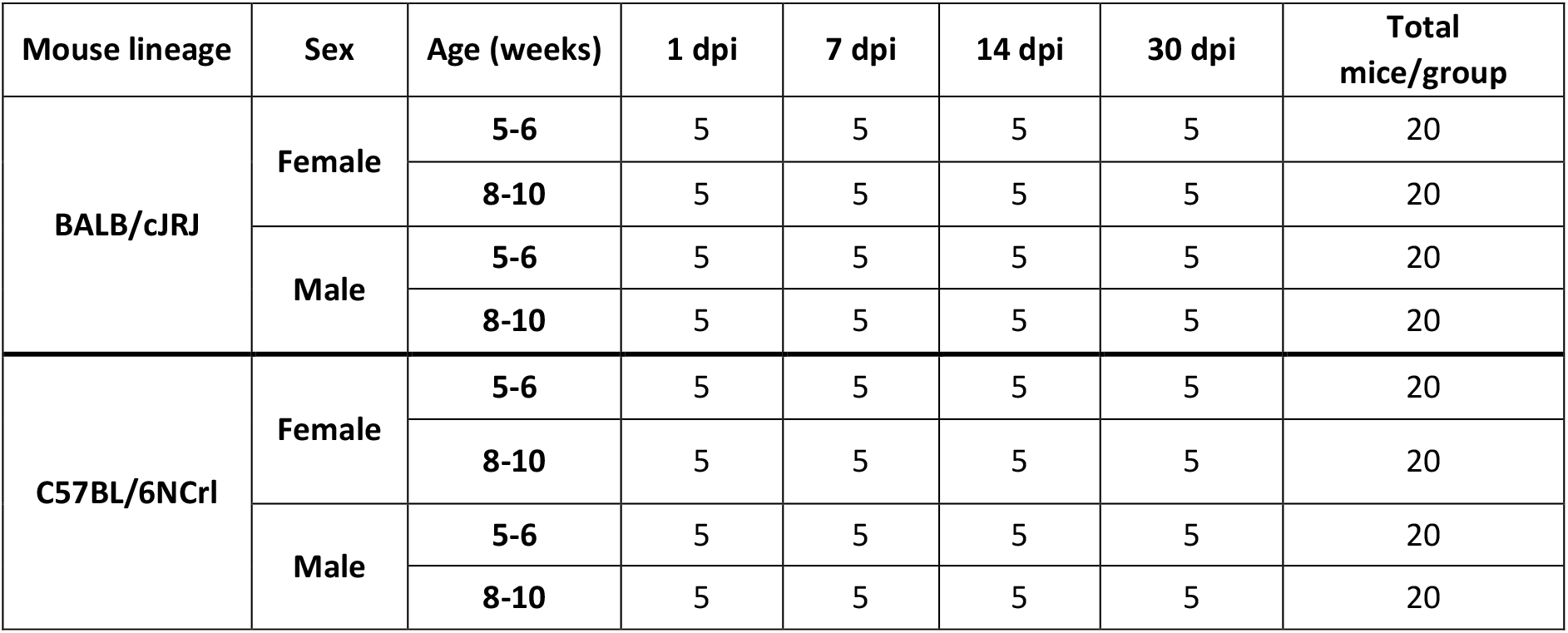
Experimental design.

